# Cellular N-myristoyl transferases Are Required for Mammarenavirus Multiplication

**DOI:** 10.1101/2024.08.01.606235

**Authors:** Haydar Witwit, Carlos Betancourt, Beatrice Cubitt, Roaa Khafaji, Heinrich Kowalski, Nathaniel Jackson, Chengjin Ye, Luis Martinez-Sobrido, Juan C. de la Torre

## Abstract

The mammarenavirus matrix Z protein plays critical roles in virus assembly and cell egress, whereas heterotrimer complexes of a stable signal peptide (SSP) together with glycoprotein subunits GP1 and GP2, generated via co-and post-translational processing of the surface glycoprotein precursor GPC, form the spikes that decorate the virion surface and mediate virus cell entry via receptor-mediated endocytosis. The Z protein and SSP undergo N-terminal myristoylation by host cell N-myristoyltransferases (NMT1 and NMT2), and G2A mutations that prevent myristoylation of Z or SSP have been shown to affect Z mediated virus budding and GP2 mediated fusion activity required to complete the virus cell entry process. In the present work, we present evidence that the validated on-target specific pan NMT inhibitor DDD85464 exerts a potent antiviral activity against the prototypic mammarenavirus lymphocytic choriomeningitis virus (LCMV) that correlated with reduced Z budding activity and GP2 mediated fusion activity, as well as proteasome mediated degradation of the Z protein. The potent anti-mammarenaviral activity of DDD85646 was also observed with the hemorrhagic fever causing mammarenaviruses Junin (JUNV) and Lassa (LASV) viruses. Our results support exploration of NMT inhibition as a broad-spectrum antiviral against human pathogenic mammarenaviruses.

## 1. Introduction

Mammarenaviruses are enveloped viruses with a bi-segmented negative-stranded (NS) RNA genome [1]. Each genome RNA segment, L (ca 7.3 kb) and S (ca 3.5 kb), uses an ambisense coding strategy to direct the synthesis of two polypeptides in opposite orientation, separated by a non-coding intergenic region (IGR). The L genome segment encodes the viral RNA dependent RNA polymerase (L), and Z matrix protein, whereas the S genome segment encodes the viral glycoprotein precursor (GPC) and the viral nucleoprotein (NP). GPC is co-translationally cleaved by signal peptidase to produce a 58 amino acid stable signal peptide (SSP) and GP that is post-translationally processed by the cellular site 1 protease (S1P) to yield the mature GP1 and GP2 subunits that together with the SSP form the spikes that decorate the virus surface and mediate cell entry via receptor-mediated endocytosis [2–5]. GP1 mediates binding to the cellular receptor and GP2 the pH-dependent fusion event in the late endosome required for release of the virus ribonucleoprotein (vRNP) complexes into the cell cytoplasm where they direct the replication and transcription of the viral genome [6,7].

Early studies shown that myristic acid analogs inhibited multiplication of the mammarenavirus Junin (JUNV) [8], and subsequently N-myristoylation was shown to be required for the role of the mammarenavirus matrix Z protein in assembly and budding [9], and for the role of the SSP in the GP2-mediate fusion event [10]. These findings were based on the use of 2-hydroxy-myristic acid (2-HMA) and 2-HMA analogs as inhibitors of N-myristoyltransferases (NMT1 and NMT2) responsible for catalyzing N-myristoylation in mammalian cells [8,9]. However, recent studies have demonstrated that 2-HMA acts off-target and does not inhibit N-myristoylation in a concentration range consistent with activity on NMT [11]. Therefore, we have revisited the contribution of N-myristoylation in mammarenavirus infection using the validated on-target specific NMT1/2 inhibitor DDD85462 [12,13]. Here, we present evidence that DDD85646 exhibits a very potent antiviral activity against the prototypic mammarenavirus lymphocytic choriomeningitis virus (LCMV) in cultured cells. Cell-based assays probing different steps of the LCMV life cycle indicated that DDD85462 interfered with Z budding activity and GP2 mediated fusion between viral and cellular membranes, a process that requires the participation of myristoylated SSP [10,14,15]. Consistent with the proposed role of a glycine-specific N-degron pathway in quality control of protein N-myristoylation [16], treatment with DDD85646 resulted in targeting of Z for degradation via the proteasome pathway, which may have also contributed to the robust DDD85646 mediated inhibition of production of infectious virus progeny.

Several mammarenaviruses cause hemorrhagic fever (HF) disease in humans and pose important public health problems in their endemic regions [17–22]. Thus, the mammarenavirus Lassa virus (LASV), the causative agent of Lassa fever (LF), is highly prevalent in Western Africa where it is estimated to infect several hundred thousand people yearly resulting in a high number of LF cases associated with high morbidity and mortality. To date, there are no FDA-approved mammarenavirus-specific therapeutics, and current LASV therapy is limited to an off-label use of ribavirin for which efficacy remains controversial [23]. Several small molecules, including the broad-spectrum mammarenavirus RNA-directed RNA polymerase inhibitor favipiravir and the mammarenavirus glycoprotein GP2-mediated fusion inhibitor ST-193, have shown promising anti-mammarenaviral effects in various animal models of mammarenavirus-induced human diseases [24–27]. Nevertheless, the identification and characterization of additional safe and effective anti-mammarenaviral drugs can facilitate the implementation of combination therapy, an approach known to counteract the emergence of drug-resistant variants often observed with mono therapy strategies. Moreover, the emergence of viral variants resistant to host-targeting antivirals (HTAs) is usually significantly reduced or entirely absent, but HTAs can be associated with significant side effects. However, side effects associated with the use of HTAs might be manageable in the case of acute infections, such as HF disease caused by mammarenaviruses, where the duration of the treatment would be rather short.

The potent anti-mammarenaviral activity of DDD85646 was also observed with other mammarenaviruses including LASV and Junin virus (JUNV), the causative agent of Argentine HF disease, suggesting that inhibition of NMT can possibly be exploited as a broad-spectrum antiviral against human pathogenic mammarenaviruses. Aberrant NMT expression has been identified in cancer cells and inhibition of myristoylation is being actively pursued as a novel anticancer treatment strategy. Accordingly, the small molecule NMT inhibitor PCLX-001 has been shown to be safe and well tolerated in humans [28,29], supporting the interest of exploring the repurposing of NMT inhibitors to treat infections by human pathogenic mammarenaviruses.

## 2. Materials and Methods

### 2.1. Cells and viruses

Grivet (Chlorocebus aethiops) Vero E6 (ATCC CRL-1586), and Homo sapiens A549 (ATCC CCL-185), and HEK 293T (ATCC CRL-3216) cell lines were maintained in Dulbecco’s modified eagle medium (DMEM) (ThermoFisher Scientific, Waltham, MA, USA) containing 10% heat-inactivated fetal bovine serum (FBS), 2 mM of L-glutamine, 100 μg/ml of streptomycin, and 100 U/ml of penicillin. HAP1 cells were obtained from Horizon Discovery and grown in IMDM supplemented with 10% FBS and 1% Pen-Strep. CRISPR/Cas9 mediated NMT1 and NMT2 knockout (KO) HAP1 cell lines have been described [12]. The recombinant LCMV expressing the green fluorescent protein (GFP) fused to LCMV NP via a P2A ribosomal skipping sequence (rLCMV/GFP-P2A-NP, referred to as rLCMV/GFP; a single cycle infectious rLCMV expressing the Zoanthus sp. green fluorescent protein (ZsG) fused to LCMV NP via a P2A ribosomal skipping sequence (rLCMVΔGPC/ZsG-P2A-NP, here referred to as rLCMVΔGPC/ZsG) [30]; the tri-segmented form of the live attenuated vaccine strain Candid#1 of JUNV expressing green fluorescent protein (GFP, r3Can/GFP) [31], the tri-segmented forms of TCRV expressing GFP (r3TCRV/GFP) [32], reassortant ML29 expressing GFP (r3ML29/GFP) [33], and the tri-segmented form of LASV Josiah strain [34] have been described. The rLCMV/Z-HA was generated using described procedures [35], but with the L segment containing a C-terminal HA-tagged version of Z.

### 2.2. Compounds

DDD85646 (2,6-Dichloro-4-[2-(1-piperazinyl)-4-pyridinyl]-N-(1,3,5-trimethyl-1H-pyrazol-4-yl)-benzenesulfonamide, CAS No. 1215010-55-1) was synthesized as described [12] [13], dissolved in DMSO at 25 mM and kept in aliquots at –20°C, and IMP-1088 ( 5-[3,4-difluoro-2-[2-(1,3,5-trimethyl-1H-pyrazol-4-yl)ethoxy]phenyl]-N,N,1-trimethyl-1H-indazole-3-methanamine, Cat No. 25366-1) was purchased from Cayman Chemical, dissolved in methyl acetate at 10 mM and kept in aliquot at -20. Clemizole Hydrochloride (MFG No. HY-30234A-5MG, MedChemExpress) was purchased from Fisher Scientific (Cat. No. 502082225). F3406-2010 was purchased from DC Chemicals (Cat. No. DC11877). Ribavirin was purchased from fSigma-Aldrich (R9644). MG132 was purchased from Sigma-Aldrich (MFG No. M7449). Sodium Citrate Buffer (0.1M, pH 5.0, Sterile) was purchased from bioWorld (MFG No. 40121003-1). FluoroBrite DMEM was purchased from Life Technologies (Cat. No. A1896701).

### 2.3. Cell cytotoxicity assay and CC_50_ determination

Cell viability was assessed using the CellTiter 96 AQueous One Solution Reagent (Promega, Madison). This method determines the number of viable cells based on conversion of formazan product from 3-(4,5-dimethylthazol-2-yl)-5-(3-carboxymethoxyphenyl)-2-(4-sulfophenyl)-2H-tetrazolim by nicotinamide adenine dinucleotide phosphate (NADPH) or nicotinamide adenine dinucleotide phosphate (NADH) generated in living cells. A549 cells were plated on 96-well clear bottom plates (4.0 x 10^4^ cells/well). Serial dilutions (2-fold) of each compound were added to cells, and at 72 h after drug treatment, CellTiter 96 AQueous One Solution Reagent was added and incubated for 35 min (37°C and 5% CO_2_). Signal was quantified using Cytation 5 reader (BioTek, Winooski, VT, USA). The resulting optical densities were normalized to the vehicle treated (DMSO) control samples, which was assigned a value of 100%. Half-maximal cytotoxic concentrations (CC_50_) were determined using GraphPad Prism, v10 (Prism10).

### 2.4. EC_50_ Determination

For EC_50_ determination, cells were seeded on 96-well clear-bottom black plates (4.0 x 10^4^ cells/well) and 20 h later infected (MOI 0.05) with rLCMV/GFP-P2A-NP. After 90 minutes adsorption, virus inoculum was aspirated off and compounds-containing media added to cells. At 72 h pi, cells were fixed with 4% paraformaldehyde, and GFP expression was determined by fluorescence using a fluorescent plate reader (Cytation 5, BioTek, Winooski, VT, USA). Mean relative fluorescence units were normalized to vehicle control (DMSO) treated cells, which were assigned a value of 100%. Half-maximal effective concentrations (EC_50_) were determined using Prism10. The selectivity index (SI) for each compound was determined using the ratio CC_50_/EC_50_.

### 2.5. Viral Growth Kinetics

For growth kinetics, virus was added to cells (500 µl/well in a M12-well plate) at the indicated MOI. After 90 min of adsorption (37°C and 5% CO_2_) virus inocula were removed, cells were washed once with DMEM 2% FBS, and fresh media containing the indicated compounds and concentrations were added and infection allowed to proceed at 37°C and 5% CO_2_. At the indicated times post-infection (pi), cell-culture supernatants (CCS) were collected, and viral titers were determined using a focus-forming assay (FFA) [36].

### 2.6. LCMV minigenome assay

The LCMV minigenome (MG) assay was performed as described [37]. Briefly, HEK 293T cells were cultured on poly-L-lysine-treated M-12 well plates (4.5 x 10^5^ cells/well). Cells were transfected using lipofectamine 2000 (2.5 µl/µg of DNA) (ThermoFisher Scientific) with Pol II-based expression plasmids (pCAGGS) for T7 RNA polymerase (pC-T7, 0.5 µg), NP (pC-NP, 0.3 µg) and L (pC-L, 0.3 µg) together with a plasmid directing intracellular synthesis of an LCMV MG expressing the chloramphenicol acetyl transferase (CAT) reporter gene under a T7 promoter (pT7-MG/CAT, 0.5 µg). After 5 h, transfection mixture was replaced with fresh media and incubated for 72 h at 37°C and 5% CO_2_. At 72 h post-transfection, whole cell lysates were harvested to determine protein expression levels of CAT using CAT ELISA kit (Roche, Sydney, Australia). Briefly, whole cell lysates were prepared with 0.5 ml of lysis buffer, and 10 µl of each sample were used for the reaction. Diluted samples were added onto CAT ELISA plates and incubated for 1 h at 37°C. After incubation with samples, plates were washed, and primary antibody (anti-CAT-digoxigenin) and secondary antibody (anti-CAT-digoxigenin-peroxidase) were added sequentially followed by the substrate. After 20 min, absorbance was measured using the ELISA reader at 405 nm for samples and 490 nm for the reference.

### 2.7. Budding assay

The luciferase-based budding assay was performed as described [38]. Briefly, HEK 293T cells were seeded on poly-L-lysine-coated M12 well plates (3.5 x 10^5^ cells/well). After overnight incubation, 2 µg of DNA of either pC-LCMV-Z-Gaussia luciferase (GLuc) or pC-LCMV-mutant Z[G2A]-GLuc, or pC-LASV-Z was transfected using Lipofectamine 2000 (2.5 µl/µg of DNA).

After 5 h, transfection mixtures were replaced with fresh media containing the indicated compounds. After 48 h, cell culture supernatant (CCS) containing virion-like particles (VLPs) were harvested and clarified by low-speed centrifugation to remove cell debris. Aliquots (20 µl each) from CCS samples were added to 96-well black plates (VWR, West Chester, PA, USA), and 50 µl of SteadyGlo luciferase reagent (Promega) was added to each well. Whole cell lysates (WCL) from the same samples were processed to determine cell-associated activity of GLuc. Luminescence signal was measured using the Berthold Centro LB 960 luminometer (Berthold Technologies, Oak Ridge, TN, USA). Activity (relative light units) of GLuc in CCS and WCL were used as surrogates of Z expression. Budding efficiency was defined as the ratio Z_VLP_/Z_VLP_+Z_WCL_.

### 2.8. Western blotting

Whole cell lysates were prepared in lysis buffer (250 mM NaCl, 50 mM Tris-HCl [pH 7.5], 0.5% Triton X-100, 10% glycerol). Samples were denatured for 5 minutes at 95°C. Then, 12.5 μg of each sample were separated by SDS-PAGE using stain-free gel (Bio-Rad, Hercules, CA, USA), transferred to low fluorescence PVDF membrane (Bio-Rad, Hercules, CA, USA), and immunoblotted with anti-HA (Genescrip, Piscataway, NJ, USA). Bands were visualized with the chemiluminescent substrate (ThermoFisher Scientific).

### 2.9. Virus titration

Virus titers were determined by FFA [36]. Serial (10-fold) dilutions of samples were done in DMEM containing 2% FBS and used to infect Vero E6 cell monolayers in 96-well plates (2 x 10^4^ cells/well). At 20 h pi, cells were fixed with 4% paraformaldehyde in phosphate-buffered saline. Foci of cells infected with rLCMV/GFP were determined by epifluorescence.

### 2.10. RNA isolation and characterization

Total cellular RNA was isolated using TRI reagent (TR 118) (Molecular Research Center, Cincinnati, OH, USA) according to the manufacturer’s instructions, resuspended in sodium citrate pH 6.4 and stored at -80^0^C. RNA samples were analyzed by Northern blot hybridization or RT-qPCR.

#### Northern blotting

RNA samples (4 μg) were fractionated by 2.2 M formaldehyde-agarose (1.2%) gel electrophoresis. The gel was washed once with warm each H_2_O and 10 mM NaPO_4_, and RNA transferred in 20X SSC [3 M sodium chloride, 0.3 M sodium citrate] to a Magnagraph membrane (NJTHYA0010, Osmonics MagnaGraph nylon) using the rapid downward transfer system (TurboBlotter). Membrane-bound RNA was cross-linked by exposure to UV light (automated 12 seconds duration, twice), the membrane was washed with MilliQ water and stained with methylene blue (MB) to reveal the 18S and 28S RNA plus RNA ladder. After image acquisition by Image Quant LAS 4000 (GE Healthcare), 1% SDS solution was used to remove MB staining, and the membrane was hybridized using QuickHyb (#201220–12 Agilent) to a ^32^P-labeled dsDNA NP. Hybridization was performed at 65°C overnight. The DNA probe was prepared according to the supplier’s protocol using a DecaPrime kit (Ambion). After overnight hybridization, the membrane was washed twice with 2X SSC–1% SDS at 65°C, followed by two washes with 0.2X SSC–0.1% SDS at 65°C, and then exposed to an X-ray film using Typhoon Trio Imager (GE Healthcare).

#### RT-qPCR

RNA (1 μg) was reverse-transcribed to cDNA using the SuperScript™ IV first-strand synthesis system (Thermo Fisher Scientific). To amplify LCMV NP and the housekeeping gene GAPDH, Powerup SYBR (A25742, Life Technologies) was used. The following primers were used for the amplification: NP forward (F): 5′ CAGAAATGTTGATGCTGGACTGC-3′, and NP reverse (R): 5′-CAGACCTTGGCTTGCTTTACACAG-3′ [39]; GAPDH F: 5′-CATGAGAAGTATGACAACAGCC-3′, and GAPDH R: 5′-TGAGTCCTTCCACGATACC-3′. ISG15 F: 5′-CAGGACGACCTGTTCTGGC-3′, and ISG15 R: 5′-GATTCATGAACACGGTGCTCAGG-3′, MX1 F: 5′-GCAGCTTCAGAAGGCCATGC-3′, MX1 R: 5′-CCTTCAGGAACTTCCGCTTGTC-3**′.**

### 2.11. GP2 mediated fusion assay

HEK293T cells were plated on poly-L-lysine-treated 24-well plates (2.5 x 10^5^ cells/well). Next day, cells were transfected with a pCAGGS plasmid expressing GFP protein (50 ng/well), along with either an empty pCAGGS plasmid or pCAGGS plasmids expressing LCMV GPC or LASV GPC proteins (1 µg/well). Lipofectamine 3000 reagent was used for transfection according to the manufacturer’s protocol. Cells were incubated with the transfection mixture for 5 hours, washed once with media, and then replaced with media with or without DDD85646 (5 µM). At 24 hours post-transfection, cells were treated with acidified (pH) 5 or neutral (pH 7.2) media for 15 minutes, followed by a wash with DMEM and returned to DMEM containing 10% FBS, and monitored over time for the appearance of syncytia using a fluorescence microscope. Cells were fixed with 4% PFA, washed with DPBS, and imaged at 20X magnification using a Keyence BZ-X710 microscope.

### 2.12. Assessment of NP to Z ratio

A549 cells were seeded onto 96-well clear bottom plates (4.0 x 10^4^ cells/well), infected with rLCMV/Z-HA (MOI 1) and treated with serial dilutions (3-fold) of the indicated compound. At 72 h after drug treatment, cells were fixed with 4% PFA. NP was stained with a rat monoclonal antibody VL4 against NP (Bio C Cell, West Lebanon, NH, USA) conjugated to Alexa Fluor 488 and Z protein was stained with an anti-HA antibody (Cat No. A01244, Genescript, USA) conjugated with Alexa Fluor 647. DNA was probed with DAPI. The resulting optical signals were normalized to vehicle (DMSO) control group, which was adjusted to 100%. Results were plotted using Prism10.

### 2.13. Immunofluorescence and subcellular localization of NP and Z

A549 cells were plated on 96-well clear bottom optical plates (4.0 x 10^4^ cells/well). Next day, cells were infected with rLCMV/Z-HA (MOI 0.05) for 90 minutes, then media was aspirated, and replaced with DDD85646 (5μM) containing media, and at 72 h after drug treatment, cells were fixed with 4% PFA. NP was identified with the rat monoclonal antibody VL4 (Bio C Cell, West Lebanon, NH, USA) conjugated to Alexa Fluor 488 and Z protein was stained with an anti-HA antibody (Cat No. A01244, Genescript, USA) conjugated with Alexa Fluor 568. DNA was probed with DAPI. HEK293T cells were plated on poly-L-Lysine 96-well clear bottom optical treated plate (2.0 x 10^4^ cells/well). Next day, cells were transfected with plasmids LCMV Z-HA, or Z-G2A-HA, and at 24 h post-transfection (h pt), media was aspirated and replaced with DDD85646 (5μM) containing media. At 48 h after drug treatment, cells were fixed with 4% PFA. Z protein was stained with an anti-HA antibody (Cat No. A01244, Genescript, USA) conjugated with Alexa Fluor 568 and DNA was probed with DAPI.

### 2.14 Statistical analyses

## 3. Results

### 3.1. Dose-dependent effect of the NMT1/2 specific inhibitor DDD85646 on LCMV multiplication

DDD85646, a validated potent and selective pan NMT inhibitor [13,40], exhibited a potent dose-dependent inhibitory effect on rLCMV/GFP-P2A-NP multiplication in A549 cells with an EC_50_ value of 0.13 μM (effective drug concentration that reduced GFP expression to 50% when compared to that of vehicle-treated controls) (Fig.1). The inhibitory effect of DDD85646 on LCMV multiplication was not a consequence of drug-induced cell toxicity, as the CC_50_ (drug concentration that reduced cell viability by 50% when compared with that of vehicle-treated cells) of DDD85646 was > 13 μM (Fig.1), resulting in an DDD85646 selectivity index (SI = CC_50_/EC_50_) of > 100.

**Figure 1.**
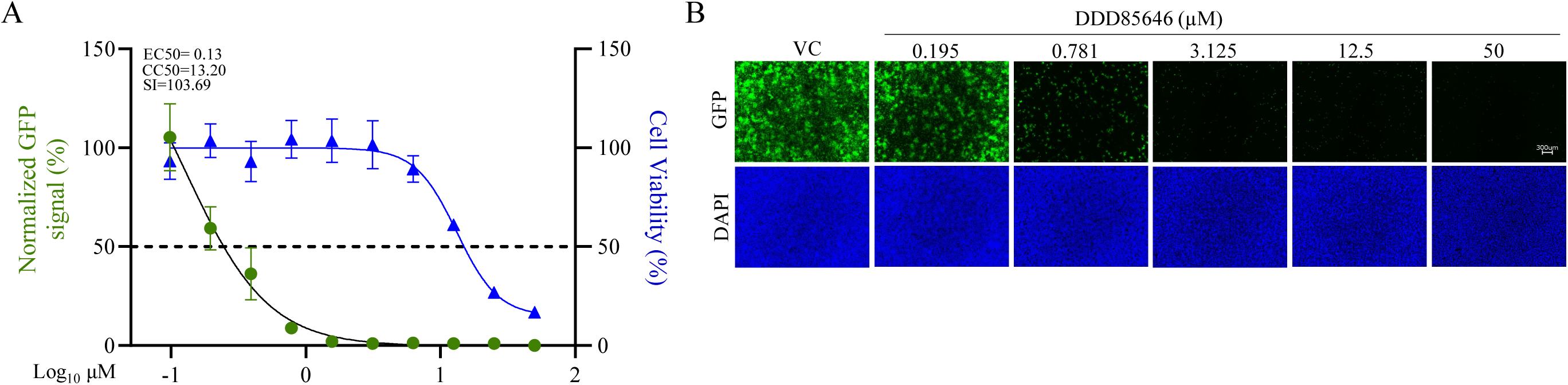
Dose-dependent effect of NMT inhibitor DDD85646 on LCMV multiplication in A549 cells. A549 cells were seeded at 4 × 10^4^ cells/well into a 96-well plate, infected with rLCMV/WT-GFP-P2A-NP (MOI 0.05) and treated with DDD85646 at the indicated concentrations. At 72 h pi, cells were fixed, and the numbers of infected cells were determined by immunofluorescence. Numbers of infected cells were normalized to vehicle control (VC) infected cells and converted to % of infected cells (Α). Total viable cell number was determined by MTS/Formazan bioreduction assay using Cytation 5 reader. Results show the percentage of infected cells (four biological replicates). EC_50_ and CC_50_ values were calculated using a variable slope (four parameters) model (Prism10). Results show the mean and SD of four biological replicates. Representative immunofluorescence images of the dose-response assay of selected doses of DDD85646 are shown (B). DAPI was used to stain the DNA (nuclei). Images were taken at 4X magnification using Keyence BZ-X710 fluorescence microscope.

### 3.2. Effect of DDD85646 on LCMV multi-step growth kinetics and peak titers

To examine the effect of DDD85646 on LCMV multi-step growth kinetics in A549 cells, we infected them with rLCMV/GFP-P2A-NP (MOI 0.05) and treated them with the indicated concentrations of DDD85646 or vehicle control (VC). At the indicated hours post-infection (h pi) we collected cell culture supernatant (CCS) samples and determined virus titers using a focus forming assay (FFA) (Fig. 2A). Treatment with DDD85646 at either 10 or 5 µM resulted in lack of detection of LCMV infectious progeny, which correlated with restricted LCMV propagation within the cell monolayer in the presence of DDD85646 (Fig. 2B, C) and reduced levels of viral RNA synthesis as determined by RT-qPCR (Fig. 2D). To assess whether treatment with DDD85646 could differentially affect levels of LCMV replication and transcription, RNA samples from rLCMV/GFP-P2A-NP infected (MOI 0.05) cells were analyzed by northern blot (Fig. 2E).

**Figure 2.**
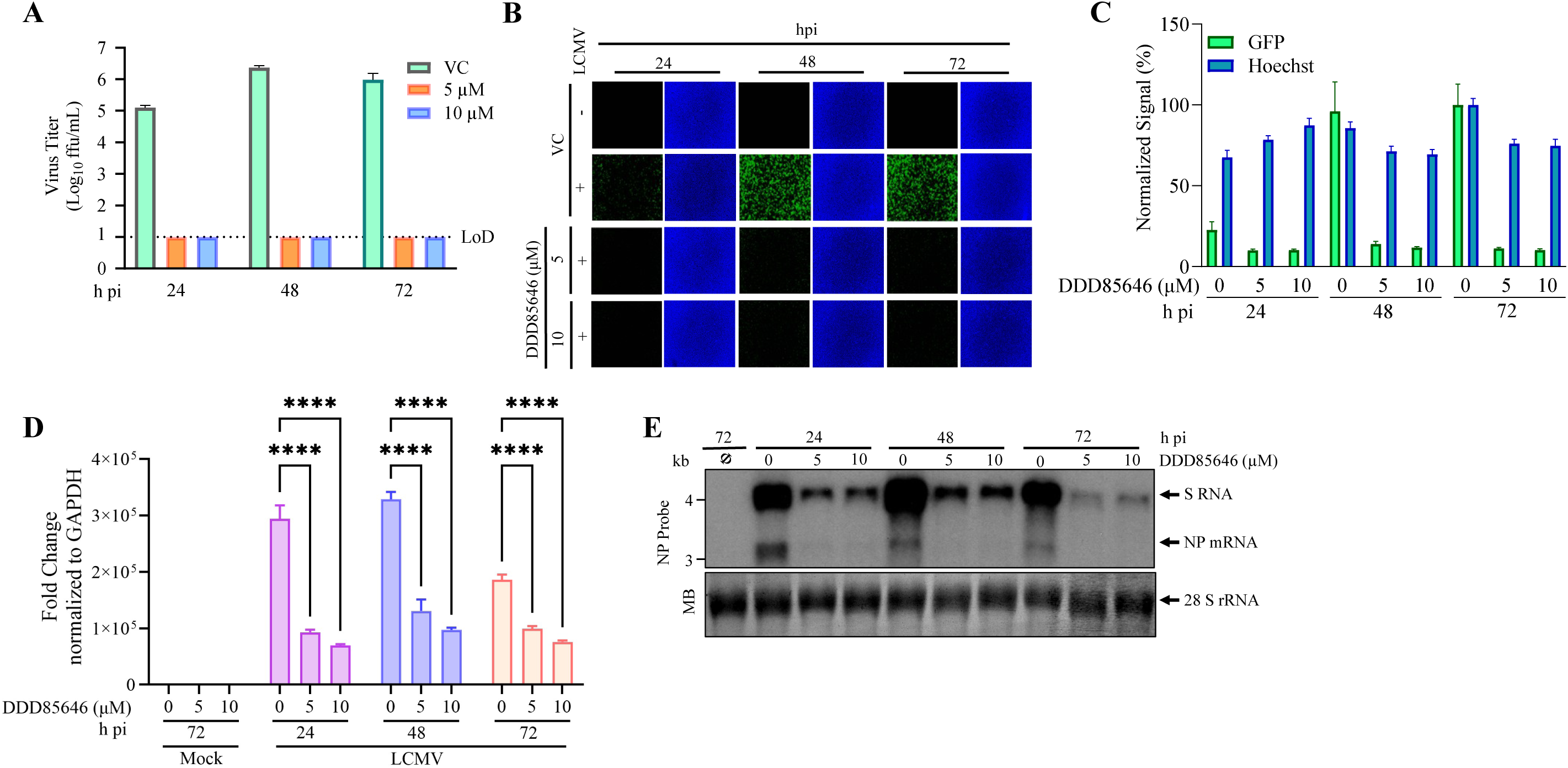
Effect of the NMT inhibitor DDD85646 on LCMV multi-step growth kinetics and peak titers in A549 cells. **A.** Effect of DDD85646 on production of infectious viral progeny. A549 cells were seeded at 2 × 10^5^ cells/well in an M12-well plate, infected with rLCMV/GFP-P2A-NP (MOI 0.05), and treated with DDD85464 (5 or 10 µM), or with VC. At indicated time points, cell culture supernatants were collected, and the titers of infectious virus were determined by focus-forming assay (FFA) using Vero E6 cells. **B, C.** Effect of DDD85646 on virus propagation. At the indicated h pi, samples from A were switched to FluoroBrite DMEM containing Hoechst dye (5 µg/mL) and stained for 15 minutes for life imaging fluorescence (B). GFP signals were determined using Cytation 5 and normalized to average of VC (72 hpi) (C). **D.** Effect of DDD85646 on viral RNA synthesis. Total cellular RNA was isolated from samples described in **B** and same RNA amount (10 ng) of each sample analyzed by RT-qPCR using random hexamers for the RT step, followed by quantitative PCR with specific primers for LCMV NP and the host cell GAPDH. The repeated measures analysis of variance with mixed effect analysis and the Dunnet’s correction for multiple comparisons were used to determine the statistical significance for the technical triplicates. Statistically significant values: **P < 0.01, ***P < 0.001, ****P < 0.0001. **E.** Effect of DDD85646 on LCMV replication and transcription. Cells were infected with rLCMV/GFP-P2A-NP (MOI 0.05), and at the indicated h pi total cellular RNA was isolated and analyzed by northern blot a LCMV NP-specific dsDNA ^32^P probe. Methylene blue (MB) staining was used to confirm similar transfer efficiency for all RNA samples.

### 3.3. Effect of DDD85646 on different steps of LCMV life cycle

To gain further insights about the mechanism by which DDD85646 exerted its antiviral activity against LCMV, we examined which steps of the virus life cycle were affected in the presence of DDD85646. To examine whether DDD85646 affected a cell entry, or post-entry, step of the LCMV life cycle we conducted a time of addition experiment using the single-cycle infectious rLCMVΔGPC/ZsG to prevent the confounding factor introduced by multiple rounds of infection (Fig. 3A). Treatment with DDD85646 starting at 1 h prior virus addition or at 2 h pi did not significantly affect the numbers of ZsG^+^ cells, whereas treatment with the inhibitor of LCMV cell entry F3406 starting at –1 h pi, but not at +2 h, resulted in over 90% reduction in the number of ZsG^+^ cells. As expected, treatment with ribavirin (Rib) (100 µM) at –1 h or +2 h pi potently reduced the number of ZsG^+^ cells detected at 48 h pi.

**Figure 3.**
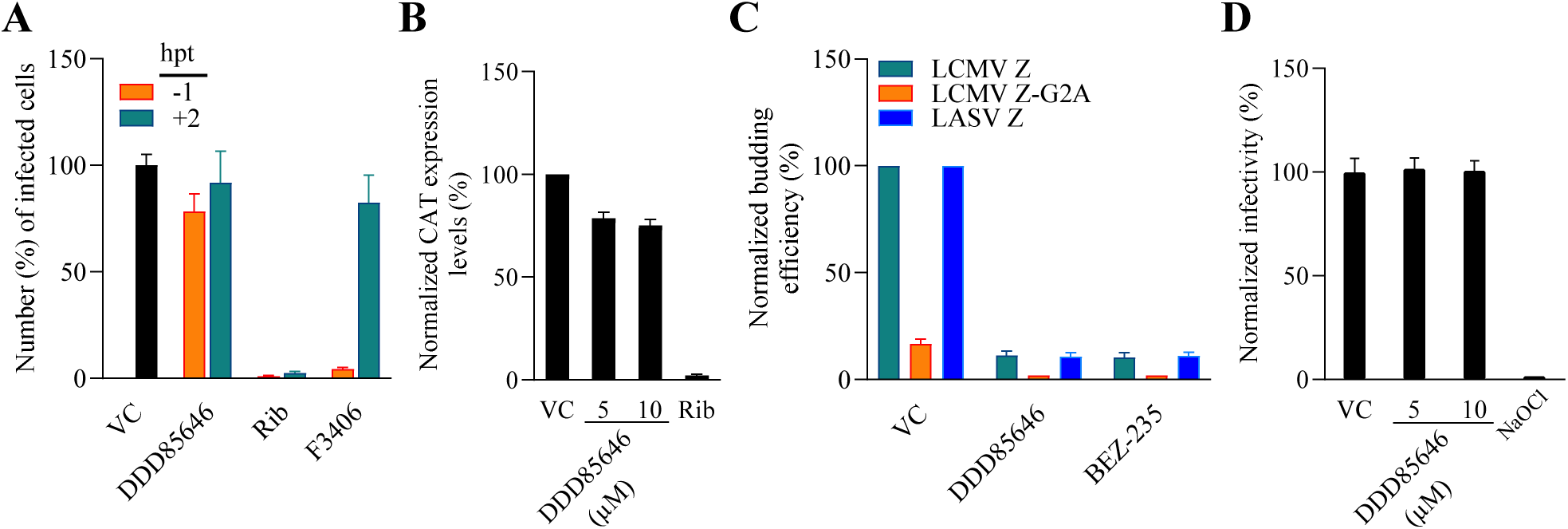
Effect of DDD85646 on different steps of the LCMV life cycle. **A.** Time of addition assay: Vero E6 cells were seeded into a 96-well plate at a density of 2 × 10^4^/well. Next day, cells were infected with the single-cycle infectious rLCMVΔGPC/ZsG (MOI 0.5) and treated with DDD85646 (5 µM), or with VC, starting 1 h before (−1 h) or after (+2 h) infection. The LCMV cell entry inhibitor F3406 (10 µM) and ribavirin (Rib) (100 µM) were used as controls. At 48 h pi, ZsG^+^ cells were assessed using Synergy™ H4 Hybrid Microplate Reader from Biotek. Values were normalized to VC-treated infected cells, the data represent an average of three biological replicates. **B.** LCMV cell-based MG assay: HEK293T cells were seeded into M24 well plate and transfected with plasmids expressing the LCMV MG-CAT, together with plasmids expressing the viral trans-acting factors NP, L polymerase, and T7 RNA polymerase, and treated with DDD85646 (5 or 10 µM), Rib and VC treated samples were used as controls. At 72 h post-transfection, cell lysates were prepared, and total protein determined using a BCA protein assay. Same amount of total protein of each sample was used to determine CAT protein expression levels using a CAT ELISA kit (Roche, Sydney, Australia) and normalized by assigning the value of 100% activity to vehicle control treated samples. **C.** Z budding activity assay: HEK293T cells were seeded onto poly-L-lysine-coated M12 wells at 1.75 x 10^5^ cells/ well. Next day, cells were transfected with either pC-LCMV-Z-GLuc, pC-LASV-Z-GLuc or pC-LCMV-Z-G2A-GLuc. At 5 h post-transfection, cells were washed three times and fed with fresh medium containing the indicated drugs and concentrations. At 48 h post-transfection, cell culture supernatant (CCS) samples were collected and whole-cell lysates (WCL) prepared. GLuc activity was determined in CCS and WCL using SteadyGlo Luciferase Pierce: Gaussia Luciferase Glow assay kit using a Berthold Centro LB 960 luminometer (Berthold Technologies, Oak Ridge, TN, USA). Activity (relative light units) of GLuc in CCS and WCL were used as surrogates of Z expression and budding efficiency defined as the ratio Z_VLP_/Z_VLP_+Z_WCL_. Budding efficiency values were normalized by assigning the value of 100% to vehicle control treated sample and plotted using Prism10. *P < 0.05, **P < 0.01, ***P < 0.001, ****P < 0.0001. **D.** Virucidal assay: 10^5^ FFU of rLCMV/GFP-P2A-NP were incubated for 30 min at RT in the presence of 0, 5, or 10 µM DDD85646, or in the presence of the validated virucidal compound sodium hypochlorite (0.5%). After treatment, samples were diluted 1000-fold in DMEM/2%FBS, resulting in concentrations (5 nM and 10 nM) that do not have noticeable anti-LCMV activity, and numbers of infectious particles determined by FFA.

To study the effect of DDD85646 on viral RNA synthesis directed by the LCMV vRNP, we examined the effect of DDD85646 on the activity of a cell based LCMV minigenome (MG) system (Fig. 3B). This MG system recapitulates LCMV RNA synthesis using an intracellular reconstituted LCMV vRNP expressing the CAT reporter gene. Reconstitution of LCMV vRNP requires co-expression of the LCMV L and NP proteins, as well as the LCMV MG vRNA. Expression levels of the MG directed CAT expression served as a comprehensive measurement of LCMV MG replication, transcription, and translation of the LCMV MG CAT reporter. Compared to vehicle-treated controls, treatment with DDD85646 at 5 or 10 µM had a modest effect on CAT expression, whereas treatment with Rib (100 µM) resulted in very low levels of CAT expression.

The mammarenavirus matrix protein Z has been shown to be the main driving force of virion budding [41]. To assess whether DDD85646 affected the Z-mediated budding process we used a published cell-based Z budding assay where the activity of the Gaussia luciferase (Gluc) reporter gene serves as a surrogate of Z budding activity [38]. We transfected HEK293T cells with a plasmid expressing LCMV Z-GLuc and treated them with DDD85646 (5 µM) or vehicle control and 48 h later measured levels of GLuc activity associated with virus-like-particles (VLPs) present in CCS, and in whole cell lysates (WCLs) (Fig. 3C). Z budding efficiency (in %) was determined by the ratio VLP-associated GLuc levels (Z_VLP_) and total GLuc levels (Z_VLP_ + Z_WCL_) times 100. BEZ-235, a known inhibitor of the Z budding activity, was used (10 µM) as a control [42]. DDD85646 had a strong inhibitory effect on LCMV and LASV Z-mediated budding.

We also examined whether DDD85646 exerted any virucidal activity on infectious LCMV virions. Treatment of LCMV infectious virions (10^5^ FFU) for 30 min at 37^0^C with DD85646 at either 5 µM or 10 µM concentrations at which DDD85646 treatment reduced production of infections LCMV progeny by > 5 logs (Fig. 2B), did not significantly affect virion infectivity, whereas treatment with 0.5% NaOCl under the same conditions resulted in a complete loss of infectivity (Fig. 3D).

### 3.4. Effect of DDD85646 on GP2 mediated fusion

Mammarenaviruses enter cells via receptor-mediated endocytosis [43–45]. In the acidic environment of the endosome, GP2 mediates a pH-dependent fusion event between viral and cellular membranes that results in the release of the vRNP into the cytoplasm of the cell where virus replication and gene transcription take place. Myristoylation of SSP has been implicated in GP2-mediated fusion [46], and therefore treatment with DDD85646 would be expected to interfere with GP2-mediated fusion. To examine this possibility, we transfected HEK293T cells with plasmids expressing LCMV (pC-LCMV-GPC) or LASV (pC-LASV-GPC) GPC, or empty plasmid (pC-E) as control, together with a plasmid expressing GFP (pC-GFP). At 5 h post-transfection cells were treated with DDD85646 (5 µM), or vehicle control (VC), next day, cells were exposed to acidic (pH 5), or neutral (pH 7.2), medium for 15 min, followed by returning cells to regular medium (pH 7.2) and cell fusion monitored over time (Fig. 4). Cells transfected with plasmids expressing LCMV or LASV GPC and exposed to pH 5 exhibited very strong fusion activity as determined by syncytial formation revealed by the pattern of GFP expression. In contrast, treatment with DDD85646 resulted in abrogation of GP2 mediated fusion upon exposure to low pH.

**Figure 4.**
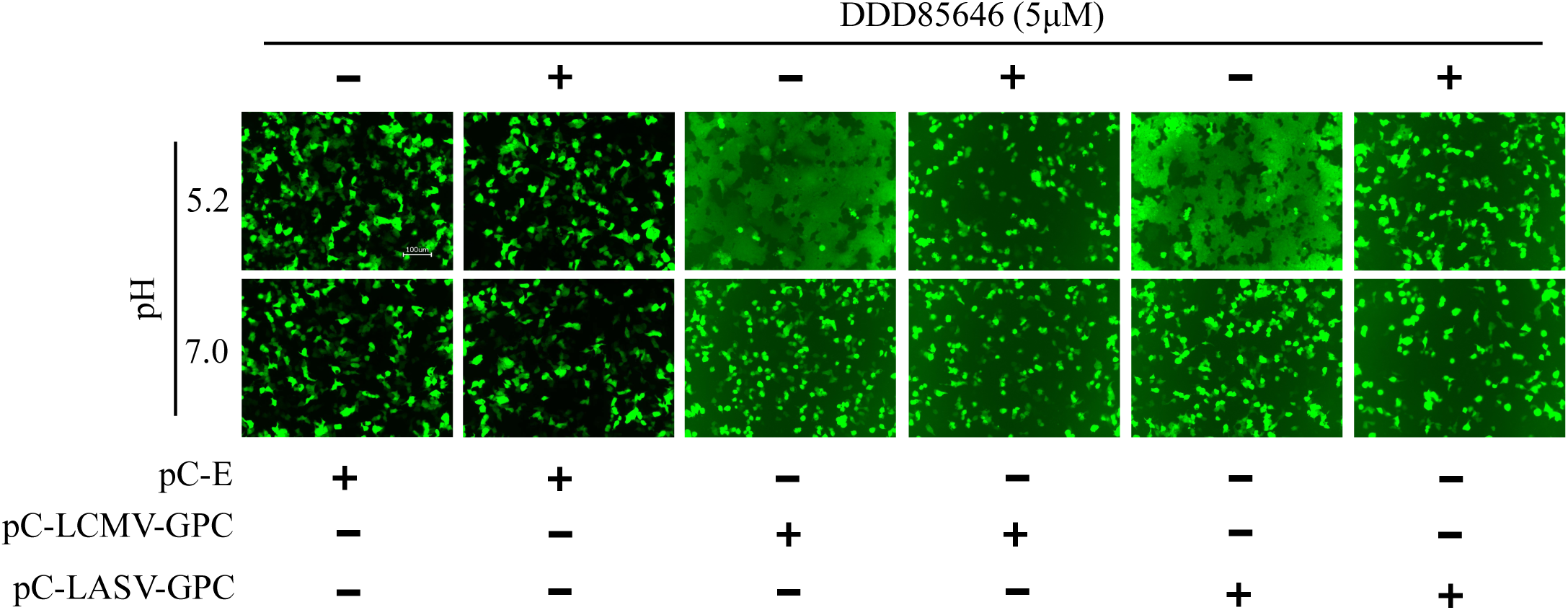
Effect of DDD85646 on LCMV and LASV GP2 mediated fusion. HEK293T cells were seeded in M24 plate at a density of 2.5 x 10^5^ cells/well, next day cells were transfected with pC-LCMV-GPC, pC-LASV-GPC or pC-E (1 μg/well), all conditions received pC-GFP (50 ng/well), at 5 h post-transfection, cells were washed once with DMEM/10% FBS and treated with DDD85646 or VC. At 24 h post-transfection cell monolayers were treated with acidic (pH 5.0) or neutral (pH 7.2) medium for 15 minutes, then return to neutral medium (DMEM/10% FBS), and examined for fusion over time. Once fusion was observed in VC treated sample, cells were fixed for IF imaging with 20X lens using Keyence BZ-X710 imaging system

### 3.5. Effect of DDD85646 on Z protein stability

N-myristoylation of the Z protein is required for its budding activity. Therefore, we expected that cell-associated levels of Z protein would increase in DDD85646, compared to vehicle, treated cells, which would account for the Z reduced budding efficiency in the presence of DDD85646. To test this hypothesis, we transfected HEK 293T cells with plasmids expressing C-terminal HA-tagged versions of Z-WT, or its mutant form Z-G2A that cannot undergo N-myristoylation. Intracellular levels of Z-WT, but not of Z-G2A, were slightly increased in the presence of DDD85646 (Fig. 5A), which was difficult to reconcile with the corresponding strong reduction in Z budding efficiency. This led us to examine whether Z protein stability was compromised in the presence of DDD85646. We found that cell-associated expression levels of Z-WT were significantly increased upon treatment with 20S proteasome inhibitor MG132 (20 μM) in the presence of DDD85646 (5 µM), supporting that as with other N-myristoylated proteins, Z expression levels is subjected to the control of the glycine-specific N-degron pathway involved in the quality control of protein N-myristoylation [16]. Consistent with this view, expression levels of Z-G2A, which does not undergo N-myristoylation, were similarly increased in the absence and presence of DDD85646 following treatment with MG132. We observed that the inhibition of the proteasome pathway by MG132 treatment resulted in increased levels of monomer and dimer forms of both WT and G2A forms of Z, but only trimers of Z-WT. Further studies, outside the scope of the present work, will examine the implications of this observation.

**Figure 5.**
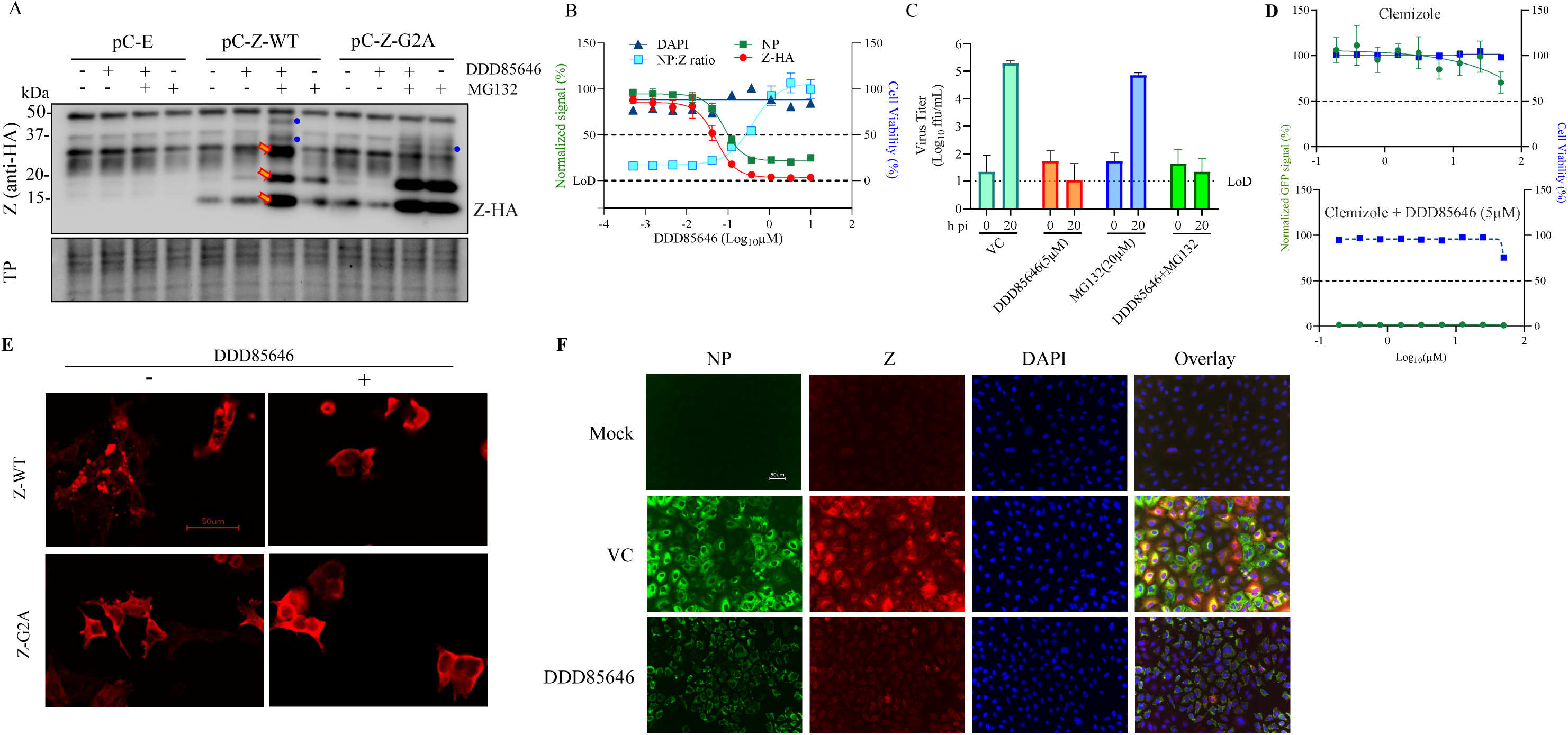
Effect of DDD85646 on Z expression levels and subcellular distribution. **A.** HEK293T cells were seeded (4x10^5^ cells/M6-well), and 16 hours later transfected with pCAGGS empty (pC-E), Z wild type (pC-Z-WT) or Z(G2A) (pC-Z-G2A). AT 24 hours post-transfection, cells were treated with DDD85646, MG132 or both, and at 24 h post-treatment, cell lysates were prepared and protein expression examined by western blot. Red arrows indicate Z protein oligomers; blue circles indicate Z protein specific associated bands. **B.** A549 cells were plated on 96-well clear bottom plates (4.0 x 10^4^ cells/well). Sixteen hours later cells were infected with rLCMV/Z-HA (MOI 1) and serial dilutions (3-fold) of DDD85646 were added to cells, started at 10μM, and at 72 h after drug treatment, cells were fixed with 4% PFA and stained for fluorescent antibodies for signal quantification, each point represents the mean of four biological replicates. **C.** A549 cells were infected (MOI = 0.1) with rLCMV/GFP-P2A-NP. After 90 minutes adsorption, the inoculum was removed, cells washed once and medium containing either MG132 (20 µM), or DD85646 (5 µM) or MG132 + DDD85646, added to cells. Cell culture supernatants were collected at 0 (after adsorption) and 24 h post-infection and virus titers determined by FFA. **D.** A549 cells were seeded at 4 × 10^4^ cells/well into a 96-well plate, infected with rLCMV/GFP-P2A-NP (MOI 0.05) and treated with incremental dose of clemizole, or clemizole in presence of 5μM of DDD85646. At 72 h pi, cells were fixed, and the numbers of infected cells were determined by IF. **E.** HEK293T cells were plated on poly-L-Lysine 96-well clear bottom optical treated plate (2.0 x 10^4^ cells/well). Next day, cells were transfected with LCMV Z-HA or Z-G2A-HA mutant, 24 h post-transfection (h pt), media was replaced with DDD85646 (5μM) contained media, and at 48 h after drug treatment, cells were fixed with 4% PFA. Images were taken using Keyence BZX-710 microscope at 20x magnification then were zoomed digitally x1.5. Images were subjected to Point-Spread Function (PSF) then deconvoluted using DeconvolutionLab2 (Fiji ImageJ) with the following setting: Iterative Constraint Tikhonov-Miller algorithm at 100 iteration N, 1 step γ, and low reg. λ. **F.** A549 cells were plated on 96-well clear bottom optical plates (2.0 x 10^4^ cells/well). Next day, cells were infected rLCMV/Z-HA (MOI 0.05), 90 minutes later, treated with DDD85646 (5μM) contained media, and at 72 h after drug treatment, cells were fixed with 4% PFA, and co-labeled with anti-HA (Z) and anti-NP antibodies, DAPI for DNA Z protein is red, NP is green, and DNA is blue, images zoomed digitally x3.

To further investigate the role of myristoylation on Z expression, we examined the effect of DDD85646 on Z expression levels in LCMV-infected cells. Because of the lack of suitable Z-specific antibodies, for this experiment we used a rLCMV expressing a C-terminal HA tagged Z (rLCMV/Z-HA). We infected A549 cells with rLCMV/Z-HA (MOI 1) and treated them with the indicated DDD85646 concentrations and at 72 h pi determined the NP:Z ratios by measuring immunofluorescent signals obtained with a mouse monoclonal antibody to HA (Z protein) and the rat monoclonal antibody VL4 to LCMV NP. Consistent with the inhibitory effect of DDD85646 on LCMV multiplication, expression levels of both Z and NP were reduced by DDD85646 treatment in a dose-dependent manner. However, the normalized NP:Z ratio was increased by DDD85646 treatment in a dose-dependent manner, supporting that Z, but not NP, was targeted for degradation upon DDD85646 mediated inhibition of myristoylation (Fig. 5B).

To assess whether proteasome mediated degradation of Z protein contributed to the antiviral activity of DDD85646 we compared the effect of MG132, in the presence and absence of DD85646, on production of LCMV infectious progeny. Treatment with MG132 did not significantly affect the inhibitory effect of DDD85646 on production of LCMV infectious progeny (Fig. 5C), and consistent with previous findings [47] MG132 did not significantly affected production of LCMV infectious progeny.

Heme oxygenase 2 (HO-2) has been shown to negatively regulate the functions of myristoylated proteins by influencing localization and trafficking of its binding partners, as well as targeting them for degradation [48,49]. Thus, pharmacological inhibition of the interaction of HO-2 with HIV-1 MA protein was shown to result in enhanced production of HIV-1 virions [48]. We therefore examined the effect of the validated HO-2 inhibitor clemizole [50–53] on LCMV multiplication using rLCMV/GFP-P2A-NP (MOI 0.05). We observed that a high (50 µM) concentration of clemizole had a rather modest effect on LCMV multiplication 72 h pi, whereas the inhibitory effect of DDD85646 (5 µM) on LCMV multiplication was not affected in the presence of clemizole (Fig. 5D).

### 3.6. Effect of DDD85646 on the subcellular distribution of Z protein

We next assessed the effect of DDD85646 treatment on the subcellular location of the Z protein. We first examined the effect of DDD85646 in HEK293T cells transfected with the pCAGGS plasmid expressing C-terminal HA-tagged versions of WT (Z-WT) or its mutant G2A (Z-G2A) Z proteins. At 48 hours post-treatment, cells were fixed and stained with an HA antibody. We observed that the Z protein exhibited a punctate appearance in the absence of DDD85646, while its distribution was homogeneous and lacked puncta in the presence of DDD85646 (5 µM). This latter distribution was also observed in cells transfected with Z-G2A and was not affected by DDD85646 treatment (Fig. 5E). Next, we examined the effect of DDD85646 on the subcellular localization of Z and NP in LCMV-infected cells. A549 cells were infected with rLCMV/Z-HA (MOI 0.05) and treated with DDD85646 (5μM), or vehicle control. Mock-infected cells served as additional control. At 48 h p.i., cells were fixed and co-labeled with anti-HA (Z) and anti-NP antibodies. Consistent with the results of studies examining the NP:Z (Fig. 5B), both NP as Z expression levels were reduced upon treatment with DDD85646, but Z exhibited a much higher degree of inhibition (Fig. 5F), further supporting that inhibition of N-myristoylation targets Z for degradation. In cells treated with DDD85646, but not with vehicle control, NP exhibited a punctate distribution (Fig. 5E), a finding that might reflect that myristoylated Z or host cell proteins can influence NP subcellular distribution [30,54–56].

### 3.7. Contribution of the type 1 interferon (T1IFN) response to DDD85646 mediated restricted LCMV multiplication

N-myristoylation has been implicated in host innate immunity defense mechanisms against microbial and viral infections [57,58]. As with many other viruses, LCMV encodes gene products that are potent viral counteracting factors of the host cell type I interferon (T1IFN) response [34,59]. We, therefore, examined whether DDD85646 induced innate immune responses that could not be counteracted by LCMV interferon counteracting factors. We used RT-qPCR to quantify levels of MX1 and ISG15 transcripts in response to DDD85646 treatment in mock- and LCMV-infected cells. Treatment with concentrations (5 and 10 µM) of DDD85646 that caused a potent inhibition of LCMV multiplication did not affect levels of MX1 or ISG15 in mock-infected cells (Fig.6). A modest, but significant, upregulation of MX1 and ISG15 mRNAs was observed in LCMV-infected cells treated with DDD85646, which likely reflects reduced levels on production of the anti-T1IFN viral factor NP [34,59].

**Figure 6.**
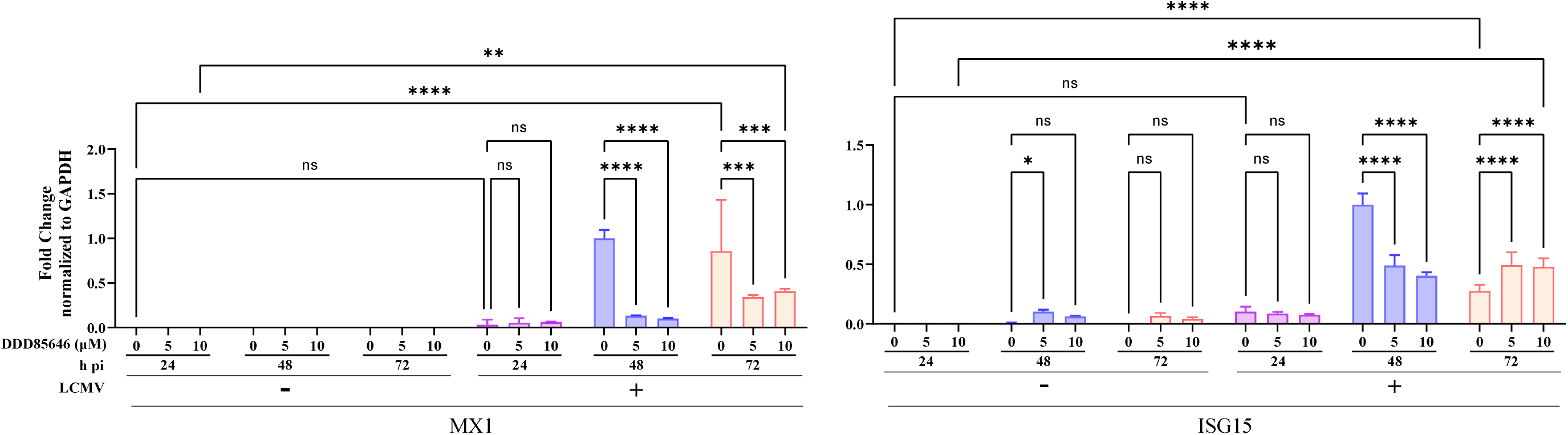
Effect of DDD85646 on expression of ISGs. A549 cells were seeded at 2 × 10^5^ cells/well in an M12-well plate, infected with LCMV/GFP- P2A-NP (MOI 0.05), and treated with DDD85464 (5 or 10 µM), or with VC. At the indicated time points, total cellular RNA was isolated and analyzed by RT-qPCR (10 ng/sample; three technical replicates). The RT step was done using random hexamers, followed by quantitative PCR with specific primers for ISGs MX1(**A**) and ISG15 (**B)**, and the housekeeping cell gene GAPDH. MX1 and ISG15 fold changes were normalized to GAPDH. The repeated measures analysis of variance with mixed effect analysis and the Dunnet’s correction for multiple comparisons were used. Statistically significant values: **P < 0.01, ***P < 0.001, ****P < 0.0001.

### 3.8. Contribution of NMT1 and NMT2 isozymes to LCMV multiplication

To investigate a possible differential role of the two ubiquitously expressed N-myristoyltransferase isozymes NMT1 and NMT2 on LCMV multiplication, we assessed multiplication of LCMV in the near-haploid fibroblast human cell line HAP1 where expression of either NMT1 or NMT2 was abrogated via CRISPR/Cas9 induced knockout (KO) lines [12]. Lack of NMT1 or NMT2 ha similar modest effect on production of LCMV infectious progeny (Fig. 7A) or cell propagation (Fig. 7B) in a multi-step growth kinetics assay. However, we observed a slight but significant differential increment at 48 hpi in NMT2-KO cells, suggesting that NMT1 might exert a dominant role on myristoylation of LCMV proteins required for normal levels of virus multiplication, a finding similar to that reported for HIV [60] and CVB3 [12].

**Figure 7.**
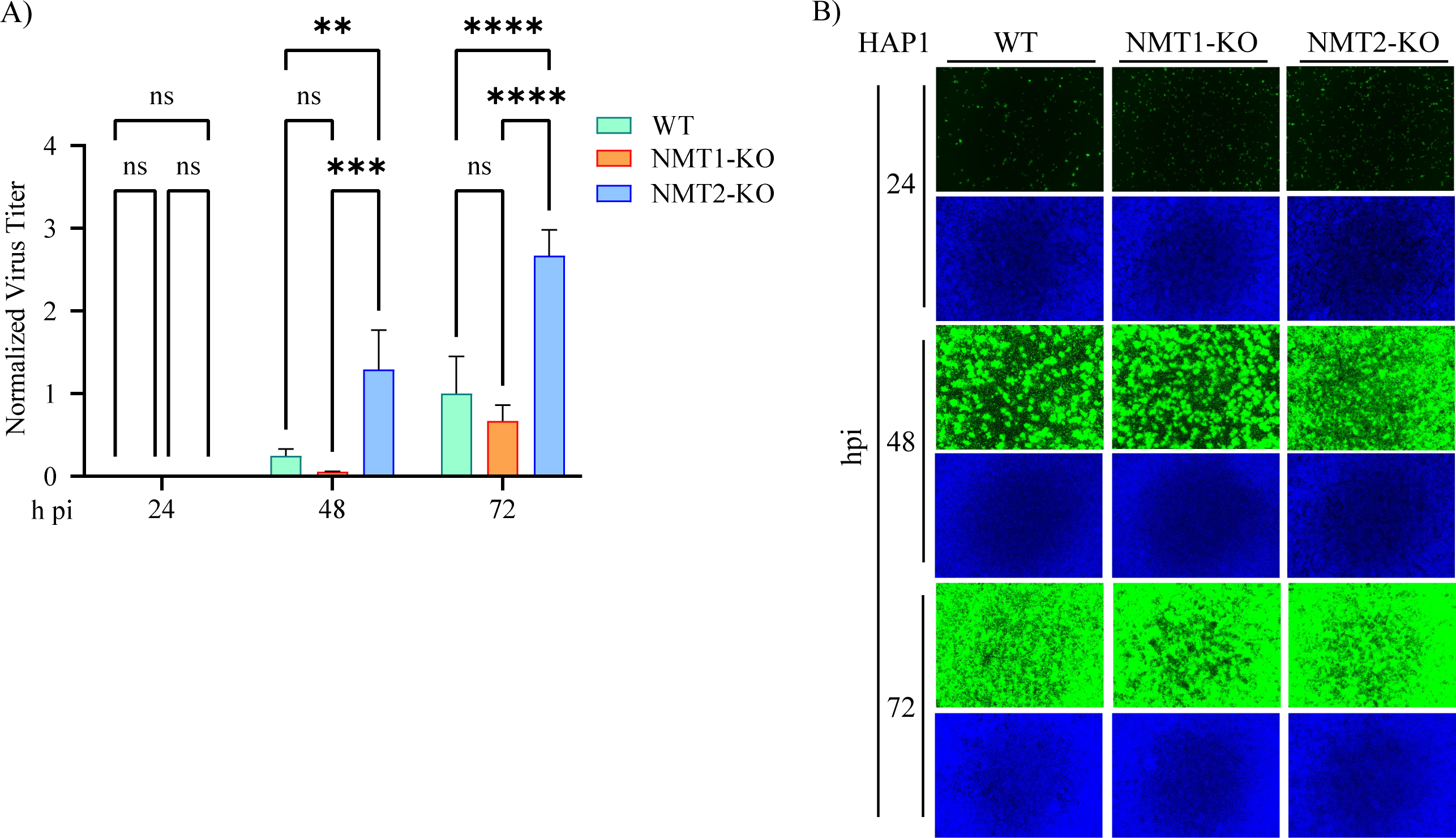
Contribution of NMT1 and NMT2 isozymes to LCMV multiplication. HAP1 NMT WT, NMT-1-KO, or NMT-2-KO cells were seeded at 2 × 10^5^ cells/well in an M12-well plate, infected with LCMV/GFP-P2A-NP (MOI 0.05). At the indicated time points, CCS were collected for virus titration. Virus titers were normalized to titers at 72 h pi in VC treated sample, which was assigned the value of 1 (**A**), and cells were fixed for immunofluorescence imaging (**B**). The repeated measures analysis of variance with mixed effect analysis and the Tukey’s correction for multiple comparisons were used. Statistically significant values: **P < 0.01, ***P < 0.001, ****P < 0.0001.

### 3.9. Effect of DDD85646 on multiplication of other mammarenaviruses

Related viruses are likely to rely on the same host cell factors for their activities, and therefore compounds targeting these host cell factors provide an opportunity for the development of broad-spectrum antiviral therapeutics. To assess whether NMT inhibitors exhibit antiviral activity against other mammarenaviruses we examined the effect of DDD85646 on multiplication of the live-attenuated vaccine strain (Candid#1) of JUNV, the LF live-attenuated vaccine candidate reassortant ML29, carrying the L segment from the non-pathogenic Mopeia virus and the S segment from LASV, and Tacaribe virus (TCRV). For these experiments we used a tri-segmented versions of Candid#1 (r3Can), ML29 (r3ML29) and TCRV (r3TCRV), expressing the GFP reporter gene. DDD85646 exhibited a potent dose-dependent inhibitory effect against these three mammarenaviruses (Fig. 8A). In contrast, and consistent with previous findings [61], Zika virus (ZIKV) multiplication in BHK21 and A549 cells was not affected by treatment with DDD85646 (Fig. 8B). DDD85646 was also effective against highly pathogenic mammarenavirus and potently inhibited multiplication in A549 cells of the HF causing mammarenavirus LASV (Fig. 8C, D).

**Figure 8.**
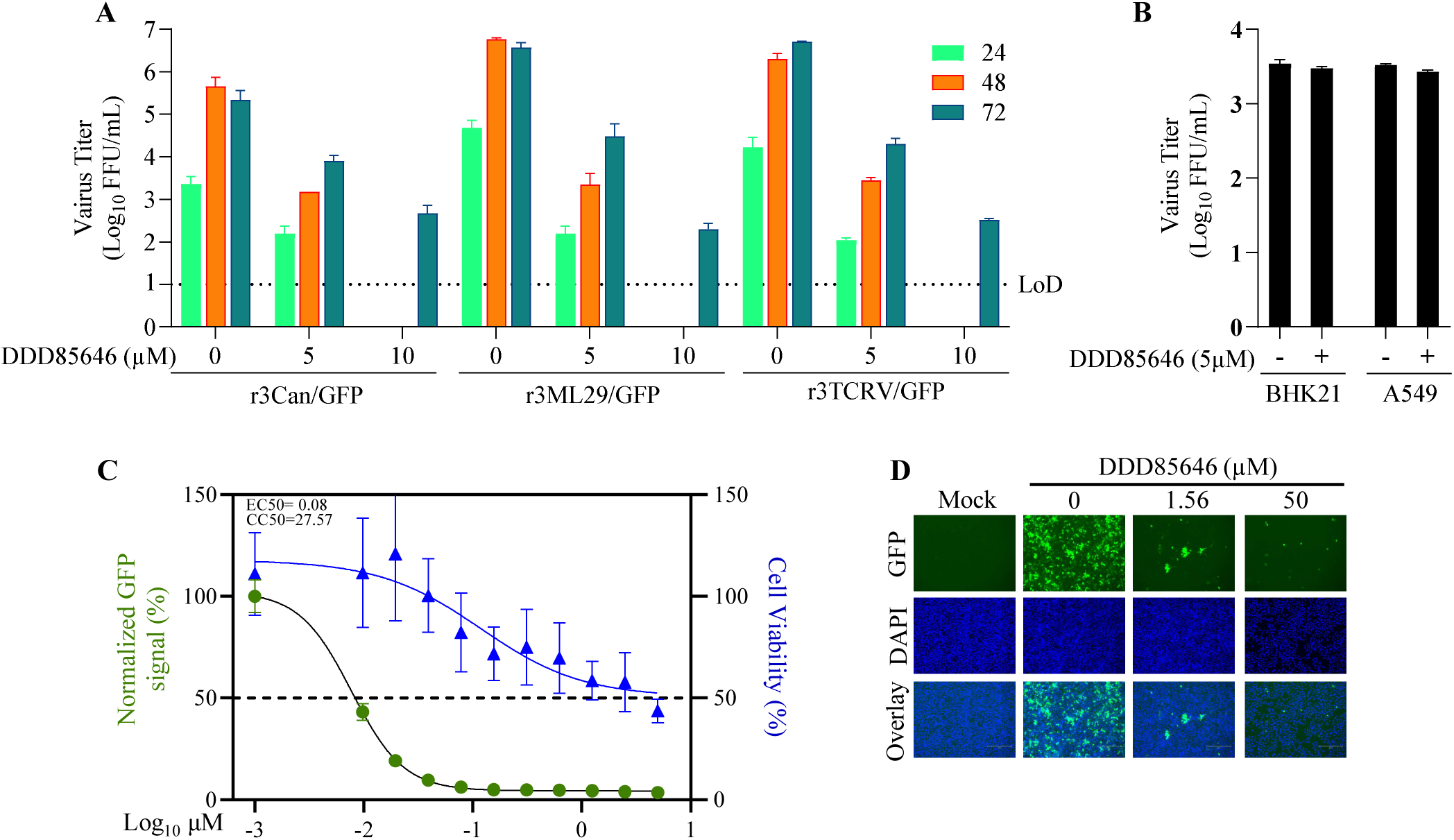
Effect of DDD85646 on multiplication of the mammarenaviruses JUNV, TCRV, ML29, and LASV, and the flavivirus ZIKV. **A.** A549 cells were infected with the indicated mammarenavirus and treated with DDD85646 (5 or 10 µM) or VC. At the indicated h pi titers of infectious virus in cell culture supernatants were determined by FFA. **B.** A549 and BHK21 cells were seeded into M24-well plate at a density of 1 x10^5^ cells/well and infected with ZIKV (MOI 0.1) for 1 hour, then treated with DDD85646 (5µM). At 24 h pi, supernatants were collected, and virus titers were determined by FFA. Titers of technical duplicates were log-transformed and plotted as mean SD (error bars). **C**. A549 cells were seeded at 4 × 10^4^ cells/well into a 96-well plate, infected with r3LASV/GFP (MOI 0.05) and treated with DDD85646 at the indicated concentrations. At 72 h pi, cells were fixed, and the numbers of infected cells were determined by immunofluorescence. Numbers of infected cells were normalized to vehicle control (VC) infected cells and converted to % of infected cells (viral infection). Cell viability was estimated based on the signal of DAPI staining. Results show the percentage of infected cells (four biological replicates). EC50 and CC50 values were calculated using a variable slope (four parameters) model (Prism10). Results show the mean and SD of four replicates. Representative immunofluorescence images of the dose-response assay of selected doses of DDD85646 are shown.

## 4. Discussion

Myristoylation of mammarenavirus SSP and Z protein plays a critical role in GP2 mediated fusion and Z mediated budding, respectively, two processes required for the completion of the mammarenavirus life cycle [1]. Myristoylation of mammalian proteins is mediated by N-myristoyltransferases (NMT) 1 and 2 [12]. Here we have presented evidence that DDD85646 [12,13], a validated specific pan-NMT inhibitor, exhibits a potent strong dose-dependent inhibitory effect on multiplication of LCMV, as well as other mammarenaviruses including the HF causing LASV and JUNV. We also observed a potent anti-LCMV and anti-LASV activity with IMP-1088, a different specific pan-NMT inhibitor [11,62] (Figure S1). DDD85646 did not affect virus cell entry and had a modest impact (20% reduction) on viral RNA synthesis mediated by vRNP in the LCMV cell-based minigenome system, whereas Z mediated budding and GP2 mediated pH-dependent fusion were strongly inhibited by DDD85646.

Myristoylation of Z is required for its budding activity, and therefore treatment with DDD85646 would be expected to result in increased levels of cell-associated Z protein. Accordingly, we observed an increase, though modest, of cell-associated Z protein following treatment with DDD85646. Levels of cell-associated Z protein in cells treated with DDD85646 markedly increased upon treatment with the proteasome inhibitor MG132, supporting that non-myristoylated Z protein is targeted for degradation via the proteasome. Z targeting for degradation was also reflected by the increased NP:Z ratios, determined by immunofluorescence, in response to treatment with increased concentrations of DDD85646. This experiment was done using a C-terminal HA-tagged version of the Z protein. However, it is highly unlikely that the HA tag influenced the outcome of the experiment, since growth kinetics and peak titers of the rLCMV-Z-HA were not significantly affected. Bioinformatic analysis revealed that HA-tagged both Z-WT and Z-G2A mutant proteins have five putative degrons [63].The functional degron region, which includes parts of the tripartite model and the tertiary structure necessary for E3 ligase engagement, is predicted to be located within 40 amino acids of the degron motif, whereas the HA tag was positioned more than 50 amino acids away from lysine 38 (K38) of the degron motif, supporting the exclusion of HA as an interfering factor. Moreover, experimental evidence supports that the HA tag does not interfere with C-terminal degrons in other proteins across different systems of eukaryotes [64–67], and we have shown that LCMV Z protein with a C-terminal HA tag is fully functional in cell-based assays virus [68]. Computational analysis of destabilizing N-terminal motifs indicated that in contrast to glycine, alanine at position 2 does not substantially contribute to protein destabilization [16]. However, Z containing the mutation G2A remained sensitive to MG132 mediated degradation, an intriguing finding that warrants studies beyond the scope of the present work.

Myristoylation of the SSP is required for GP2 mediated fusion, which could account for the inhibitory effect of DDD85646 on GP2 mediated fusion. However, we cannot rule out that proteasome mediated degradation of SSP also contributed to restricted GP2 mediated fusion in the presence of DDD85646.

The activity of myristoylated proteins can be negatively regulated by HO-2 [48]. However, the anti-LCMV activity of DDD85646 was not affected by clemizole, a validated inhibitor of HO-2 [50–53]. DDD85646 did not trigger ISGs, supporting the T1IFN-independent antiviral activity of DDD85646. Comparison of LCMV multiplication between WT, NMT1-KO, and NMT2-KO HAP1 cells indicated that NMT1 might exert a dominant role on myristoylation of LCMV proteins required for normal levels of virus multiplication. It is possible that in the absence of NMT-2, the expression of NMT1 is substantially upregulated in HAP1 cells, which could account for the more efficient production of LMCV in these cells.

Results from studies of the role of NMT in cancer cell lines support a large efficacy window between host and virus due to substantial differences in the rate of protein turnover [69].Upon initiation of treatment with an NMT inhibitor, pre-existing N-myristoylated host proteins must be degraded before the inhibitor may have an impact on the cell physiology, a process that could take several days. It should be noted that the concept of targeting a core host-lipidation process to block viral replication was validated in a phase 2A trial of an inhibitor of host prenylation of the hepatitis D virus [70]. DDD85646 exhibited a similarly potent antiviral activity against other tested mamarenaviruses including the highly pathogenic LASV, indicating its potential pan-mammarenavirus antiviral activity. The observation that mammarenaviruses have evolved to depend on host myristoylation to complete some of the steps of their life cycle suggests that this mode of action might circumvent the development of resistance to a drug that targets NMT because viral mutations would not influence the inhibitor potency against a host enzyme. Notably, a recent human phase I trial has shown that the oral, highly bioavailable, small molecule NMT inhibitor PCLX-001 is safe and well tolerated at concentrations that result in PCLX-001 plasma concentration over its EC_90_ [28]. Progress on NMT inhibitor-based cancer therapies can facilitate the repurposing of NMT inhibitors as antivirals against human pathogenic mammarenaviruses. In studies beyond the scope of the present work, we will examine the antiviral effect of PCLX-001 in a mouse model of LCMV infection.

## 5. Acknowledgments

This research was supported by NIH/NIAID grant RO1 AI142985 and R21 AI128556 to JCT. This is manuscript 30292 from The Scripps Research Institute.

---

**Figure S1.**
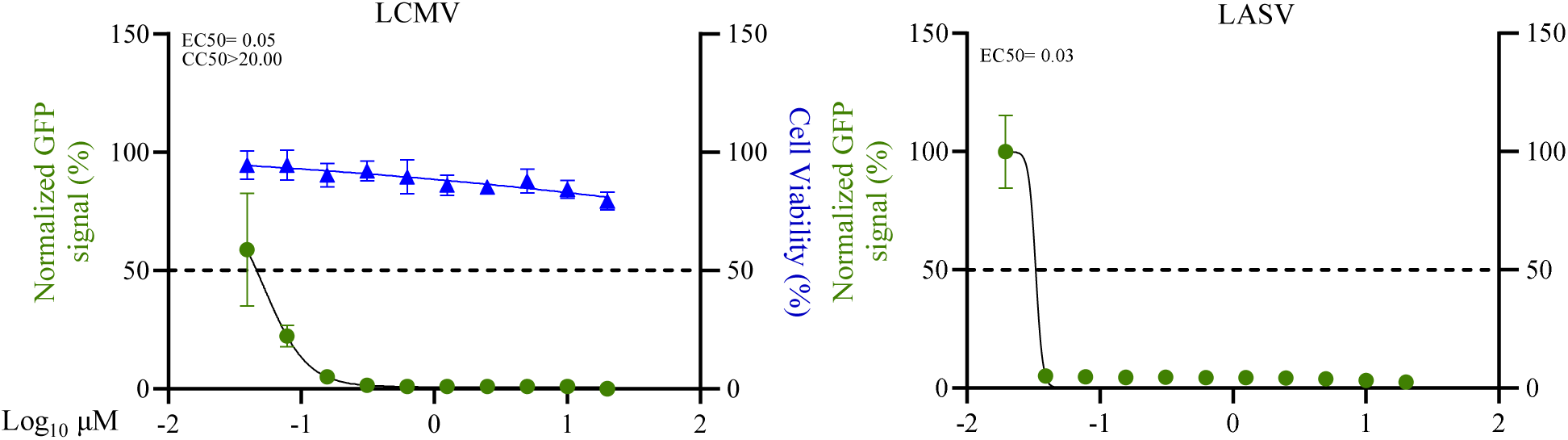
Dose-dependent effect of NMT inhibitor IMP1088 on LCMV and LASV multiplication in A549 cells. A549 cells were seeded at 4 × 10^4^ cells/well into a 96-well plate, infected with rLCMV/GFP-P2A-NP (MOI 0.05) and treated with serial dilutions (2-fold) of IMP-1088. At 72 h pi, cells were fixed, and the numbers of infected cells were determined by IF. Numbers (%) of infected cells were normalized to vehicle control (VC) infected cells. Total cell number was determined by MTS/Farmazan bioreduction assay using Cytation 5 reader. Results show the percentage of infected cells (four biological replicates). EC50 and CC50 were calculated using a variable slope (four parameters) model (Prism10).

## References

1. Radoshitzky, S.R., Buchmeier, M.J., de la Torre, J.C. 2022 Fields Virology: Emerging Viruses: Arenaviridae. Fields Virology, 7th Ed, Vol I: Emerging Viruses. Knipe, D.M., Howley, P.M., Whelan, S. (Eds). In.

2. Lenz, O.; ter Meulen, J.; Klenk, H.-D.; Seidah, N.G.; Garten, W. The Lassa Virus Glycoprotein Precursor GP-C Is Proteolytically Processed by Subtilase SKI-1/S1P. Proc Natl Acad Sci U S A 2001, 98, 12701–12705, doi:10.1073/pnas.221447598.

3. Beyer, W.R.; Pöpplau, D.; Garten, W.; von Laer, D.; Lenz, O. Endoproteolytic Processing of the Lymphocytic Choriomeningitis Virus Glycoprotein by the Subtilase SKI-1/S1P. J Virol 2003, 77, 2866–2872, doi:10.1128/JVI.77.5.2866-2872.2003.

4. Kunz, S.; Edelmann, K.H.; de la Torre, J.-C.; Gorney, R.; Oldstone, M.B.A. Mechanisms for Lymphocytic Choriomeningitis Virus Glycoprotein Cleavage, Transport, and Incorporation into Virions. Virology 2003, 314, 168–178, doi:10.1016/s0042-6822(03)00421-5.

5. Rojek, J.M.; Lee, A.M.; Nguyen, N.; Spiropoulou, C.F.; Kunz, S. Site 1 Protease Is Required for Proteolytic Processing of the Glycoproteins of the South American Hemorrhagic Fever Viruses Junin, Machupo, and Guanarito. J Virol 2008, 82, 6045–6051, doi:10.1128/JVI.02392-07.

6. Fedeli, C.; Moreno, H.; Kunz, S. Novel Insights into Cell Entry of Emerging Human Pathogenic Arenaviruses. J Mol Biol 2018, 430, 1839–1852, doi:10.1016/j.jmb.2018.04.026.

7. Hallam, S.J.; Koma, T.; Maruyama, J.; Paessler, S. Review of Mammarenavirus Biology and Replication. Front Microbiol 2018, 9, 1751, doi:10.3389/fmicb.2018.01751.

8. Cordo, S.M.; Candurra, N.A.; Damonte, E.B. Myristic Acid Analogs Are Inhibitors of Junin Virus Replication. Microbes and Infection 1999, 1, 609–614, doi:10.1016/S1286-4579(99)80060-4.

9. Perez, M.; Greenwald, D.L.; de La Torre, J.C. Myristoylation of the RING Finger Z Protein Is Essential for Arenavirus Budding. J Virol 2004, 78, 11443–11448, doi:10.1128/JVI.78.20.11443-11448.2004.

10. York, J.; Nunberg, J.H. Role of the Stable Signal Peptide of Junín Arenavirus Envelope Glycoprotein in pH-Dependent Membrane Fusion. J Virol 2006, 80, 7775–7780, doi:10.1128/JVI.00642-06.

11. Kallemeijn, W.W.; Lueg, G.A.; Faronato, M.; Hadavizadeh, K.; Goya Grocin, A.; Song, O.-R.; Howell, M.; Calado, D.P.; Tate, E.W. Validation and Invalidation of Chemical Probes for the Human N-Myristoyltransferases. Cell Chem Biol 2019, 26, 892–900.e4, doi:10.1016/j.chembiol.2019.03.006.

12. Corbic Ramljak, I.; Stanger, J.; Real-Hohn, A.; Dreier, D.; Wimmer, L.; Redlberger-Fritz, M.; Fischl, W.; Klingel, K.; Mihovilovic, M.D.; Blaas, D.;, et al. Cellular N-Myristoyltransferases Play a Crucial Picornavirus Genus-Specific Role in Viral Assembly, Virion Maturation, and Infectivity. PLoS Pathog 2018, 14, e1007203, doi:10.1371/journal.ppat.1007203.

13. Frearson, J.A.; Brand, S.; McElroy, S.P.; Cleghorn, L.A.T.; Smid, O.; Stojanovski, L.; Price, H.P.; Guther, M.L.S.; Torrie, L.S.; Robinson, D.A.;, et al. N-Myristoyltransferase Inhibitors as New Leads to Treat Sleeping Sickness. Nature 2010, 464, 728–732, doi:10.1038/nature08893.

14. York, J.; Romanowski, V.; Lu, M.; Nunberg, J.H. The Signal Peptide of the Junín Arenavirus Envelope Glycoprotein Is Myristoylated and Forms an Essential Subunit of the Mature G1-G2 Complex. J Virol 2004, 78, 10783–10792, doi:10.1128/JVI.78.19.10783-10792.2004.

15. York, J.; Nunberg, J.H. Myristoylation of the Arenavirus Envelope Glycoprotein Stable Signal Peptide Is Critical for Membrane Fusion but Dispensable for Virion Morphogenesis. J Virol 2016, 90, 8341–8350, doi:10.1128/JVI.01124-16.

16. Timms, R.T.; Zhang, Z.; Rhee, D.Y.; Harper, J.W.; Koren, I.; Elledge, S.J. A Glycine-Specific N-Degron Pathway Mediates the Quality Control of Protein N-Myristoylation. Science 2019, 365, eaaw4912, doi:10.1126/science.aaw4912.

17. Andersen, K.G.; Shapiro, B.J.; Matranga, C.B.; Sealfon, R.; Lin, A.E.; Moses, L.M.; Folarin, O.A.; Goba, A.; Odia, I.; Ehiane, P.E.;, et al. Clinical Sequencing Uncovers Origins and Evolution of Lassa Virus. Cell 2015, 162, 738–750, doi:10.1016/j.cell.2015.07.020.

18. Richmond, J.K.; Baglole, D.J. Lassa Fever: Epidemiology, Clinical Features, and Social Consequences. BMJ 2003, 327, 1271–1275.

19. Yun, N.E.; Walker, D.H. Pathogenesis of Lassa Fever. Viruses 2012, 4, 2031–2048, doi:10.3390/v4102031.

20. Radoshitzky, S.R.; Buchmeier, M.J.; de la Torre, J.C. Fields Virology: Emerging Viruses; Knipe, D.M., Howley, P.M., Griffin, D., Lamb, R., Martin, M., Roizman, B., Straus, S., Eds.; Arenaviridae: The Viruses and Their Replication.; 5th ed.; Lippincott Williams & Wilkins (LWW), 2007; Vol. II;

21. Bray, M. Pathogenesis of Viral Hemorrhagic Fever. Curr Opin Immunol 2005, 17, 399– 403, doi:10.1016/j.coi.2005.05.001.

22. Grant, A.; Seregin, A.; Huang, C.; Kolokoltsova, O.; Brasier, A.; Peters, C.;Paessler, S. Junín Virus Pathogenesis and Virus Replication. Viruses 2012, 4, 2317–2339, doi:10.3390/v4102317.

23. Salam, A.P.; Cheng, V.; Edwards, T.; Olliaro, P.; Sterne, J.; Horby, P. Time to Reconsider the Role of Ribavirin in Lassa Fever. PLOS Neglected Tropical Diseases 2021, 15, e0009522, doi:10.1371/journal.pntd.0009522.

24. Gowen, B.B.; Juelich, T.L.; Sefing, E.J.; Brasel, T.; Smith, J.K.; Zhang, L.; Tigabu, B.; Hill, T.E.; Yun, T.; Pietzsch, C.;, et al. Favipiravir (T-705) Inhibits Junín Virus Infection and Reduces Mortality in a Guinea Pig Model of Argentine Hemorrhagic Fever. PLoS Negl Trop Dis 2013, 7, e2614, doi:10.1371/journal.pntd.0002614.

25. Safronetz, D.; Rosenke, K.; Westover, J.B.; Martellaro, C.; Okumura, A.; Furuta, Y.; Geisbert, J.; Saturday, G.; Komeno, T.; Geisbert, T.W.;, et al. The Broad-Spectrum Antiviral Favipiravir Protects Guinea Pigs from Lethal Lassa Virus Infection Post-Disease Onset. Sci Rep 2015, 5, 14775, doi:10.1038/srep14775.

26. Mendenhall, M.; Russell, A.; Smee, D.F.; Hall, J.O.; Skirpstunas, R.; Furuta, Y.; Gowen, B.B. Effective Oral Favipiravir (T-705) Therapy Initiated after the Onset of Clinical Disease in a Model of Arenavirus Hemorrhagic Fever. PLOS Neglected Tropical Diseases 2011, 5, e1342, doi:10.1371/journal.pntd.0001342.

27. Cashman, K.A.; Smith, M.A.; Twenhafel, N.A.; Larson, R.A.; Jones, K.F.; Allen, R.D.; Dai, D.; Chinsangaram, J.; Bolken, T.C.; Hruby, D.E.;, et al. Evaluation of Lassa Antiviral Compound ST-193 in a Guinea Pig Model. Antiviral Res 2011, 90, 70–79, doi:10.1016/j.antiviral.2011.02.012.

28. Sangha, R.S.; Jamal, R.; Spratlin, J.L.; Kuruvilla, J.; Sehn, L.H.; Weickert, M.; Berthiaume, L.G.; Mackey, J.R. A First-in-Human, Open-Label, Phase I Trial of Daily Oral PCLX-001, an NMT Inhibitor, in Patients with Relapsed/Refractory B-Cell Lymphomas and Advanced Solid Tumors. JCO 2023, 41, e15094–e15094, doi:10.1200/JCO.2023.41.16_suppl.e15094.

29. Sangha, R.; Davies, N.M.; Namdar, A.; Chu, M.; Spratlin, J.; Beauchamp, E.; Berthiaume, L.G.; Mackey, J.R. Novel, First-in-Human, Oral PCLX-001 Treatment in a Patient with Relapsed Diffuse Large B-Cell Lymphoma. Curr Oncol 2022, 29, 1939–1946, doi:10.3390/curroncol29030158.

30. Iwasaki, M.; Minder, P.; Caì, Y.; Kuhn, J.H.; Yates, J.R.; Torbett, B.E.; de la Torre, J.C. Interactome Analysis of the Lymphocytic Choriomeningitis Virus Nucleoprotein in Infected Cells Reveals ATPase Na+/K+ Transporting Subunit Alpha 1 and Prohibitin as Host-Cell Factors Involved in the Life Cycle of Mammarenaviruses. PLoS Pathog 2018, 14, e1006892, doi:10.1371/journal.ppat.1006892.

31. Emonet, S.F.; Seregin, A.V.; Yun, N.E.; Poussard, A.L.; Walker, A.G.; de la Torre, J.C.; Paessler, S. Rescue from Cloned cDNAs and In Vivo Characterization of Recombinant Pathogenic Romero and Live-Attenuated Candid #1 Strains of Junin Virus, the Causative Agent of Argentine Hemorrhagic Fever Disease. J Virol 2011, 85, 1473–1483, doi:10.1128/JVI.02102-10.

32. Ye, C.; de la Torre, J.C.; Martínez-Sobrido, L. Development of Reverse Genetics for the Prototype New World Mammarenavirus Tacaribe Virus. J Virol 2020, 94, e01014–20, doi:10.1128/JVI.01014-20.

33. Lukashevich, I.S.; Paessler, S.; de la Torre, J.C. Lassa Virus Diversity and Feasibility for Universal Prophylactic Vaccine. F1000Res 2019, 8, F1000 Faculty Rev-134, doi:10.12688/f1000research.16989.1.

34. Witwit, H.; Khafaji, R.; Salaniwal, A.; Kim, A.S.; Cubitt, B.; Jackson, N.; Ye, C.; Weiss, S.R.; Martinez-Sobrido, L.; de la Torre, J.C. Activation of Protein Kinase Receptor (PKR) Plays a pro-Viral Role in Mammarenavirus-Infected Cells. Journal of Virology 2024, 98, e01883–23, doi:10.1128/jvi.01883-23.

35. Sánchez, A.B.; de la Torre, J.C. Rescue of the Prototypic Arenavirus LCMV Entirely from Plasmid. Virology 2006, 350, 370–380, doi:10.1016/j.virol.2006.01.012.

36. Battegay, M.; Cooper, S.; Althage, A.; Bänziger, J.; Hengartner, H.; Zinkernagel, R.M. Quantification of Lymphocytic Choriomeningitis Virus with an Immunological Focus Assay in 24- or 96-Well Plates. J Virol Methods 1991, 33, 191–198, doi:10.1016/0166-0934(91)90018-u.

37. Perez, M.; de la Torre, J.C. Characterization of the Genomic Promoter of the Prototypic Arenavirus Lymphocytic Choriomeningitis Virus. J Virol 2003, 77, 1184–1194, doi:10.1128/JVI.77.2.1184-1194.2003.

38. Capul, A.A.; de la Torre, J.C. A Cell-Based Luciferase Assay Amenable to High-Throughput Screening of Inhibitors of Arenavirus Budding. Virology 2008, 382, 107–114, doi:10.1016/j.virol.2008.09.008.

39. McCausland, M.M.; Crotty, S. Quantitative PCR (QPCR) Technique for Detecting Lymphocytic Choriomeningitis Virus (LCMV) in Vivo. J Virol Methods 2008, 147, 167– 176, doi:10.1016/j.jviromet.2007.08.025.

40. Paape, D.; Bell, A.S.; Heal, W.P.; Hutton, J.A.; Leatherbarrow, R.J.; Tate, E.W.; Smith, D.F. Using a Non-Image-Based Medium-Throughput Assay for Screening Compounds Targeting N-Myristoylation in Intracellular Leishmania Amastigotes. PLoS Negl Trop Dis 2014, 8, e3363, doi:10.1371/journal.pntd.0003363.

41. Perez, M.; Craven, R.C.; de la Torre, J.C. The Small RING Finger Protein Z Drives Arenavirus Budding: Implications for Antiviral Strategies. Proc Natl Acad Sci U S A 2003, 100, 12978–12983, doi:10.1073/pnas.2133782100.

42. Urata, S.; Ngo, N.; de la Torre, J.C. The PI3K/Akt Pathway Contributes to Arenavirus Budding. J Virol 2012, 86, 4578–4585, doi:10.1128/JVI.06604-11.

43. Borrow, P.; Oldstone, M.B. Mechanism of Lymphocytic Choriomeningitis Virus Entry into Cells. Virology 1994, 198, 1–9, doi:10.1006/viro.1994.1001.

44. Castilla, V.; Mersich, S.E.; Candurra, N.A.; Damonte, E.B. The Entry of Junin Virus into Vero Cells. Arch Virol 1994, 136, 363–374, doi:10.1007/BF01321064.

45. Radoshitzky, S.R.; Abraham, J.; Spiropoulou, C.F.; Kuhn, J.H.; Nguyen, D.; Li, W.; Nagel, J.; Schmidt, P.J.; Nunberg, J.H.; Andrews, N.C.;, et al. Transferrin Receptor 1 Is a Cellular Receptor for New World Haemorrhagic Fever Arenaviruses. Nature 2007, 446, 92–96, doi:10.1038/nature05539.

46. York, J.; Romanowski, V.; Lu, M.; Nunberg, J.H. The Signal Peptide of the Junín Arenavirus Envelope Glycoprotein Is Myristoylated and Forms an Essential Subunit of the Mature G1-G2 Complex. J Virol 2004, 78, 10783–10792, doi:10.1128/JVI.78.19.10783-10792.2004.

47. Ma, X.-Z.; Bartczak, A.; Zhang, J.; Khattar, R.; Chen, L.; Liu, M.F.; Edwards, A.; Levy, G.; McGilvray, I.D. Proteasome Inhibition In Vivo Promotes Survival in a Lethal Murine Model of Severe Acute Respiratory Syndrome. Journal of Virology 2010, 84, 12419– 12428, doi:10.1128/jvi.01219-10.

48. Zhu, Y.; Luo, S.; Sabo, Y.; Wang, C.; Tong, L.; Goff, S.P. Heme Oxygenase 2 Binds Myristate to Regulate Retrovirus Assembly and TLR4 Signaling. Cell Host Microbe 2017, 21, 220–230, doi:10.1016/j.chom.2017.01.002.

49. Wang, B.; Dai, T.; Sun, W.; Wei, Y.; Ren, J.; Zhang, L.; Zhang, M.; Zhou, F. Protein N-Myristoylation: Functions and Mechanisms in Control of Innate Immunity. Cell Mol Immunol 2021, 18, 878–888, doi:10.1038/s41423-021-00663-2.

50. Vlahakis, J.Z.; Vukomanovic, D.; Nakatsu, K.; Szarek, W.A. Selective Inhibition of Heme Oxygenase-2 Activity by Analogs of 1-(4-Chlorobenzyl)-2-(Pyrrolidin-1-Ylmethyl)-1H-Benzimidazole (Clemizole): Exploration of the Effects of Substituents at the N-1 Position. Bioorganic & Medicinal Chemistry 2013, 21, 6788–6795, doi:10.1016/j.bmc.2013.07.050.

51. Floresta, G.; Fallica, A.N.; Romeo, G.; Sorrenti, V.; Salerno, L.; Rescifina, A.; Pittalà, V. Identification of a Potent Heme Oxygenase-2 (HO-2) Inhibitor by Targeting the Secondary Hydrophobic Pocket of the HO-2 Western Region. Bioorg Chem 2020, 104, 104310, doi:10.1016/j.bioorg.2020.104310.

52. Kong, X.; Vukomanovic, D.; Nakatsu, K.; Szarek, W.A. Structure-Activity Relationships of 1,2-Disubstituted Benzimidazoles: Selective Inhibition of Heme Oxygenase-2 Activity. ChemMedChem 2015, 10, 1435–1441, doi:10.1002/cmdc.201500128.

53. Intagliata, S.; Salerno, L.; Ciaffaglione, V.; Leonardi, C.; Fallica, A.N.; Carota, G.; Amata, E.; Marrazzo, A.; Pittalà, V.; Romeo, G. Heme Oxygenase-2 (HO-2) as a Therapeutic Target: Activators and Inhibitors. Eur J Med Chem 2019, 183, 111703, doi:10.1016/j.ejmech.2019.111703.

54. Thinon, E.; Serwa, R.A.; Broncel, M.; Brannigan, J.A.; Brassat, U.; Wright, M.H.; Heal, W.P.; Wilkinson, A.J.; Mann, D.J.; Tate, E.W. Global Profiling of Co- and Post-Translationally N-Myristoylated Proteomes in Human Cells. Nat Commun 2014, 5, 4919, doi:10.1038/ncomms5919.

55. Tsumagari, K.; Isobe, Y.; Ishihama, Y.; Seita, J.; Arita, M.; Imami, K. Application of Liquid-Liquid Extraction for N-Terminal Myristoylation Proteomics. Molecular & Cellular Proteomics 2023, 22, doi:10.1016/j.mcpro.2023.100677.

56. Takamitsu, E.; Otsuka, M.; Haebara, T.; Yano, M.; Matsuzaki, K.; Kobuchi, H.; Moriya, K.; Utsumi, T. Identification of Human N-Myristoylated Proteins from Human Complementary DNA Resources by Cell-Free and Cellular Metabolic Labeling Analyses. PLoS One 2015, 10, e0136360, doi:10.1371/journal.pone.0136360.

57. Jia, M.; Wang, Y.; Wang, J.; Qin, D.; Wang, M.; Chai, L.; Fu, Y.; Zhao, C.; Gao, C.; Jia, J.;, et al. Myristic Acid as a Checkpoint to Regulate STING-Dependent Autophagy and Interferon Responses by Promoting N-Myristoylation. Nat Commun 2023, 14, 660, doi:10.1038/s41467-023-36332-3.

58. Barr, S.D.; Smiley, J.R.; Bushman, F.D. The Interferon Response Inhibits HIV Particle Production by Induction of TRIM22. PLoS Pathog 2008, 4, e1000007, doi:10.1371/journal.ppat.1000007.

59. Martínez-Sobrido, L.; Emonet, S.; Giannakas, P.; Cubitt, B.; García-Sastre, A.; de la Torre, J.C. Identification of Amino Acid Residues Critical for the Anti-Interferon Activity of the Nucleoprotein of the Prototypic Arenavirus Lymphocytic Choriomeningitis Virus. J Virol 2009, 83, 11330–11340, doi:10.1128/JVI.00763-09.

60. Takamune, N.; Gota, K.; Misumi, S.; Tanaka, K.; Okinaka, S.; Shoji, S. HIV-1 Production Is Specifically Associated with Human NMT1 Long Form in Human NMT Isozymes. Microbes and Infection 2008, 10, 143–150, doi:10.1016/j.micinf.2007.10.015.

61. Immerheiser, M.; Zimniak, M.; Hilpert, H.; Geiger, N.; König, E.-M.; Bodem, J. Towards a Broad-Spectrum Antiviral, the Myristoyltransferase Inhibitor IMP-1088 Suppresses Viral Replication – the Yellow Fever NS5 Is Myristoylated 2021, 2021.03.09.434547.

62. Mousnier, A.; Bell, A.S.; Swieboda, D.P.; Morales-Sanfrutos, J.; Pérez-Dorado, I.; Brannigan, J.A.; Newman, J.; Ritzefeld, M.; Hutton, J.A.; Guedán, A.;, et al. Fragment-Derived Inhibitors of Human N-Myristoyltransferase Block Capsid Assembly and Replication of the Common Cold Virus. Nat Chem 2018, 10, 599–606, doi:10.1038/s41557-018-0039-2.

63. Szulc, N.A.; Stefaniak, F.; Piechota, M.; Soszyńska, A.; Piórkowska, G.; Cappannini, A.; Bujnicki, J.M.; Maniaci, C.; Pokrzywa, W. DEGRONOPEDIA: A Web Server for Proteome-Wide Inspection of Degrons. Nucleic Acids Research 2024, 52, W221–W232, doi:10.1093/nar/gkae238.

64. Hickey, C.M.; Breckel, C.; Zhang, M.; Theune, W.C.; Hochstrasser, M. Protein Quality Control Degron-Containing Substrates Are Differentially Targeted in the Cytoplasm and Nucleus by Ubiquitin Ligases. Genetics 2020, 217, 1–19, doi:10.1093/genetics/iyaa031.

65. Veits, G.K.; Henderson, C.S.; Vogelaar, A.; Eron, S.J.; Lee, L.; Hart, A.; Deibler, R.W.; Baddour, J.; Elam, W.A.; Agafonov, R.V.;, et al. Development of an AchillesTAG Degradation System and Its Application to Control CAR-T Activity. Current Research in Chemical Biology 2021, 1, 100010, doi:10.1016/j.crchbi.2021.100010.

66. Yeh, C.; Huang, W.; Hsu, P.; Yeh, K.; Wang, L.; Hsu, P.W.; Lin, H.; Chen, Y.; Chen, S.; Yeang, C.;, et al. The C-degron Pathway Eliminates Mislocalized Proteins and Products of Deubiquitinating Enzymes. The EMBO Journal 2021, 40, e105846, doi:10.15252/embj.2020105846.

67. Damhofer, H.; Radzisheuskaya, A.; Helin, K. Generation of Locus-Specific Degradable Tag Knock-Ins in Mouse and Human Cell Lines. STAR Protocols 2021, 2, 100575, doi:10.1016/j.xpro.2021.100575.

68. Cornu, T.I.; de la Torre, J.C. RING Finger Z Protein of Lymphocytic Choriomeningitis Virus (LCMV) Inhibits Transcription and RNA Replication of an LCMV S-Segment Minigenome. J Virol 2001, 75, 9415–9426, doi:10.1128/JVI.75.19.9415-9426.2001.

69. Thinon, E.; Morales-Sanfrutos, J.; Mann, D.J.; Tate, E.W. N-Myristoyltransferase Inhibition Induces ER-Stress, Cell Cycle Arrest, and Apoptosis in Cancer Cells. ACS Chem. Biol. 2016, 11, 2165–2176, doi:10.1021/acschembio.6b00371.

70. Koh, C.; Canini, L.; Dahari, H.; Zhao, X.; Uprichard, S.L.; Haynes-Williams, V.; Winters, M.A.; Subramanya, G.; Cooper, S.L.; Pinto, P.;, et al. Oral Prenylation Inhibition with Lonafarnib in Chronic Hepatitis D Infection: A Proof-of-Concept Randomised, Double-Blind, Placebo-Controlled Phase 2A Trial. Lancet Infect Dis 2015, 15, 1167–1174, doi:10.1016/S1473-3099(15)00074-2.

